# Functional dissection of TADs reveals non-essential and instructive roles in regulating gene expression

**DOI:** 10.1101/566562

**Authors:** Alexandra Despang, Robert Schöpflin, Martin Franke, Salaheddine Ali, Ivana Jerkovic, Christina Paliou, Wing-Lee Chan, Bernd Timmermann, Lars Wittler, Martin Vingron, Stefan Mundlos, Daniel M. Ibrahim

## Abstract

The genome is organized in megabase-sized three-dimensional units, called Topologically Associated Domains (TADs), that are separated by boundaries. TADs bring distant cis-regulatory elements into proximity, a process dependent on the cooperative action of cohesin and the DNA binding factor CTCF. Surprisingly, genome-wide depletion of CTCF has little effect on transcription, yet structural variations affecting TADs have been shown to cause gene misexpression and congenital disease. Here, we investigate TAD function *in vivo* in mice by systematically editing components of TAD organization at the *Sox9*/*Kcnj* locus. We find that TADs are formed by a redundant system of CTCF sites requiring the removal of all major sites within the TAD and at the boundary for two neighboring TADs to fuse. TAD fusion resulted in leakage of regulatory activity from the *Sox9* to the *Kcnj* TAD, but no major changes in gene expression. This indicates that TAD structures provide robustness and precision, but are not essential for developmental gene regulation. Gene misexpression and resulting disease phenotypes, however, were attained by re-directing regulatory activity through inversions and/or the re-positioning of boundaries. Thus, efficient re-wiring of enhancer promoter interaction and aberrant disease causing gene activation is not induced by a mere loss of insulation but requires the re-direction of contacts.

## Introduction

Chromosomes are organized in a specific three dimensional (3D) structure in the nuclear space, a phenomenon that is directly linked to gene regulation reviewed in^1^; ^2, 3^. On a gene locus-level, this organization is characterized by regions of high interaction called Topologically Associating Domains (TADs) that are separated from each other by so-called boundaries ^4, 5^. TAD bring distant cis-regulatory elements such as promoters and enhancers into proximity whereas boundaries are thought to act as insulators to preclude inappropriate enhancer-promoter interactions with neighboring genes or regulatory elements. This concept provides a basic framework for long range gene regulation ^6^ and also has important implications for the interpretation of genomic rearrangements (structural variations) reviewed in ^7^.

One key component for TAD and boundary formation is the zinc finger transcription factor CTCF, which acts in concert with the multi-subunit protein complex Cohesin ^8^. In the currently prevailing model, TAD formation is the result of a loop extrusion process in which cohesin molecules extrude a chromatin loop and thereby bring distant DNA fragments into spatial proximity ^9, 10^. In this model CTCF binding sites act as a barrier for the extrusion machinery in an orientation-dependent manner. This view is supported by the finding that a large fraction of TAD boundaries harbor clusters of CTCF binding sites that are characteristically positioned in divergent orientation ^3, 9^.

The importance of the CTCF/cohesin machinery for higher order chromatin architecture has further been corroborated by experimental approaches that allow for the temporary genome-wide depletion of CTCF or various subunits of the cohesin complex, circumventing their absolute requirement for cell survival ^8, 11–14^. Cells in which CTCF or cohesin is depleted loose most of their TAD structures. In spite of this dramatic loss in 3D genome organization, however, only modest effects on gene expression were observed ^8, 11, 14^. Less than half of the regulated genes exhibit elevated expression, suggesting only spurious gains in enhancer-promoter interactions in the absence of TADs and boundaries. These results are in contrast to previous findings in which the rearrangement of TADs and their boundaries were shown to have dramatic effects on gene regulation resulting in congenital disease or cancer ^15–17^. The basis for this apparent discrepancy remains unclear, raising the question about the functional importance of TADs for gene regulation and the proposed molecular pathology of SVs.

Here, we dissect the role of CTCF and TAD architecture for gene regulation in a developmental *in vivo* setting in mice. We created a series of genome-engineered mice with targeted mutations at the *Sox9*/*Kcnj* locus and analyzed their effect on 3D chromatin architecture, gene regulation, and the phenotype. The *Kcnj* and *Sox9* TADs are separated by a strong boundary, but a fusion of the TADs, as indicated by HiC, was only achieved after removal of all major CTCF sites at the boundary and within the TAD. TAD fusion, however, was not accompanied by major gene regulatory effects, suggesting that long-range gene regulation does not exclusively rely on intact TAD structures. In contrast, inversions and the insertion of boundary elements were able to re-direct regulatory activity inducing enhancer-promoter rewiring, gene misexpression and developmental phenotypes. Thus, TADs, and in particular CTCF sites are not essential for correct developmental gene expression but they can induce gene misexpression when re-directed.

## Results

### Two TADs define the regulatory landscape at the *Sox9* and *Kcnj2* locus

*Sox9* and *Kcnj2* are two adjacent genes with distinct expression patterns in the developing limb bud that are separated by a 1.7 Mb gene desert (Fig. 1). In E12.5 limb buds *Sox9* is expressed in the cartilage anlagen of the developing limbs, whereas *Kcnj2* is only weakly expressed in the distal zeugopod (Fig. 1b). Capture HiC (cHiC) from mouse limb buds shows that the locus is divided in two TADs, one harboring *Sox9* and the other *Kcnj2* and *Kcnj16* ^15^. The TAD boundary is characterized by two pairs of divergent CTCF binding sites with divergent orientation, showing strong loop formation with their neighboring boundaries (Fig. 1). Within the *Sox9-*TAD a nested substructure with various loops is linked to at least four additional CTCF binding sites (C1 to C4). To profile the regulatory landscape of the locus in more detail, we performed ATAC-seq and H3K27ac ChIP-seq from E12.5 limb buds and identified many putative enhancer regions carrying active enhancer marks (Fig. 1a). To capture the cis-regulatory activity of the locus we investigated regulatory sensors (β*globin* minimal promoter with *LacZ* reporter gene), integrated at various positions within the *Kcnj* and *Sox9* TADs (Fig. 1b). LacZ staining of E12.5 embryos revealed that all sensors within the *Sox9* TAD recapitulated the *Sox9* expression pattern, whereas sensors integrated within the *Kcnj* TAD reflected the endogenous expression pattern of *Kcnj2*. This shows that the 3D genome organization in the two TADs corresponds with the regulatory domains of the *Kcnj2* and *Sox9* genes.

**Figure 1:**
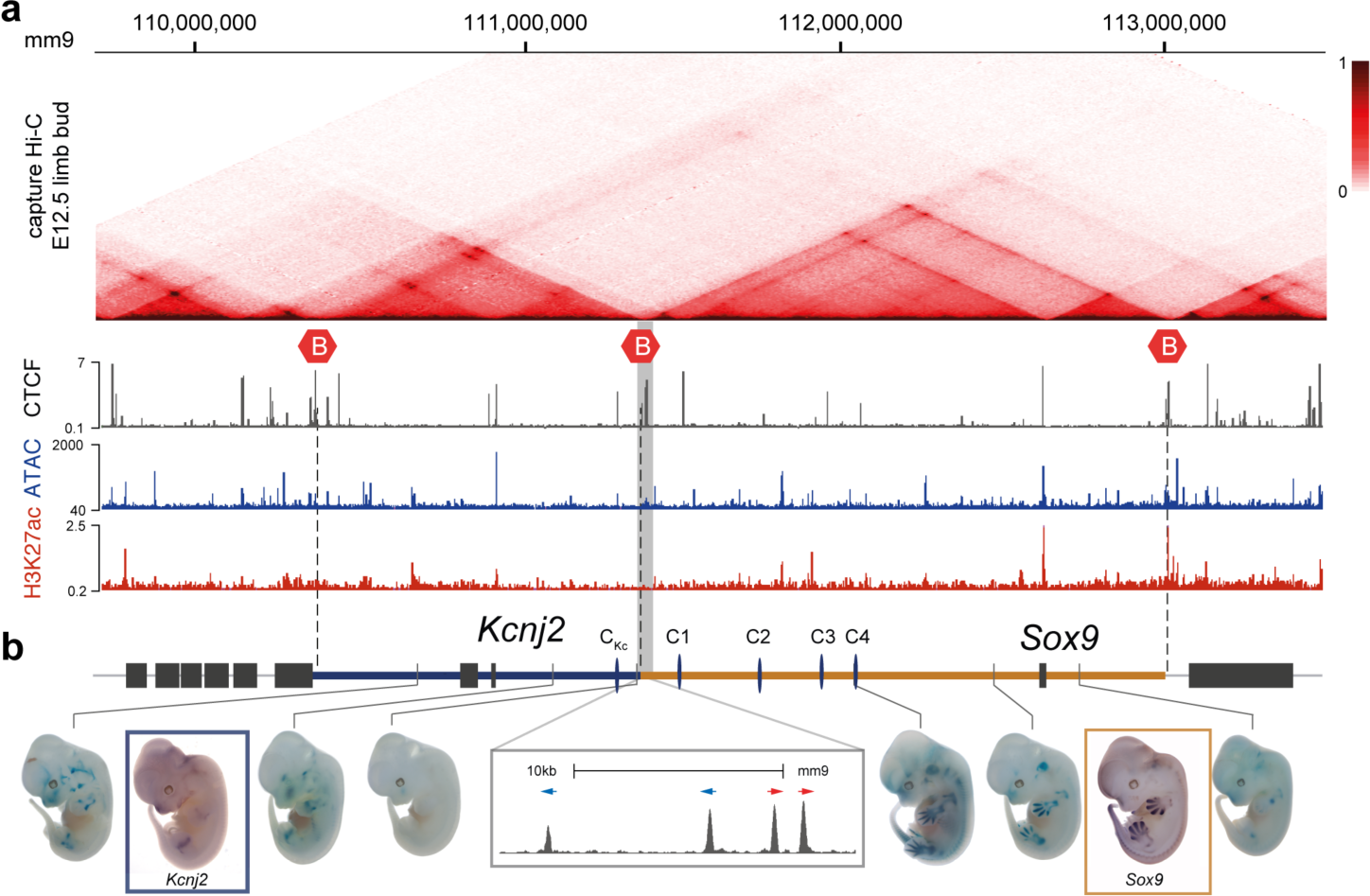
TAD configuration and regulatory activity at the *Sox9*-Locus. (A) Capture HiC from E12.5 mouse limb buds with boundaries indicated by red hexagons. CTCF ChIP-seq, ATAC-seq and H3K27ac shown below. Note multiple CTCF sites at boundaries and within TADs as well as potential cis-regulatory elements indicated by ATAC-seq tracks and H3K27ac-ChIP-seq peaks. (B) Schematic of the locus. Genes are indicated by black bars, TADs of *Kcnj2* (blue) and *Sox9* (orange). Boundary region between *Sox9* and *Kcnj*-TADs is highlighted in grey, magnification below shows the cluster of 4 divergent CTCF sites within a 15 kb region. Other major CTCF-binding sites are indicated and labeled as C1, C2, C3, C4 and C_Kc_. Lower panel shows the activity of regulatory sensors in E12.5 embryos inserted at indicated positions. Expression pattern (*WISH*)of *Kcnj2* and *Sox9* is shown for comparison.

### Boundaries and internal CTCF sites act cooperatively to form TADs

To investigate the role of CTCF in maintaining TAD structure at this locus, we produced a series of alleles in which the 4 CTCF sites at the TAD boundary were deleted, followed by a consecutive deletion of five further sites within the TADs. cHiC was performed from E12.5 limb buds to visualize the effects on TAD architecture and to quantify the contacts within the TADs (intra-TAD) and between the TADs (inter-TAD). We also produced virtual 4C interaction profiles from the cHiC data to assess contact changes of *Sox9* and *Kcnj2* in these alleles, (see Methods).

Deletion of the 4 CTCF sites that constitute the *Kcnj*/*Sox9*-TAD boundary (*ΔBor*) resulted in a moderate increase of contacts between TADs, but the two TADs remained largely separate (Fig. 2a, b) ^15^. To test whether the intra-TAD CTCF sites contribute to TAD formation we sequentially deleted, in addition to the boundary, one (*ΔBorC1*), two, and all 4 (*ΔBorC1-4*) of the major CTCF sites within the *Kcnj*/*Sox9*-TAD. Deletion of the C1 CTCF site together with the boundary deletion (*ΔBorC1*) led to a marked increase in contacts between the *Sox9-* and *Kcnj*-TADs, which became even more pronounced by the additional deletion of the C2 CTCF site (*ΔBorC1-2*) (Fig. S1). Deletion of all 4 CTCF sites (*ΔBorC1-4*) led to a further increase in inter-TAD contacts and a near complete fusion between the *Sox9* and *Kcnj-*TADs (Fig. 2c, Fig. 3a). Moreover, while the internal TAD structure and loops disappeared, new interactions between the *Sox9* and *Kcnj2* promoters, as well as between the outer TAD boundaries, emerged. Further deletion of the remaining single major CTCF site in the *Kcnj*-TAD (*ΔCTCF*) abolished all major CTCF binding sites between the *Sox9* and *Kcnj2* promoters. cHi-C from these animals showed a further increase of inter-TAD contacts between the former *Sox9* and *Kcnj-TAD*s (Fig. 2d, Fig. 3a). Taken together, our data show that the TAD boundary deletion alone does not result in TAD fusion. Instead, TAD formation and integrity is established by the TAD boundary in combination with the CTCF-mediated TAD substructure.

**Figure 2.**
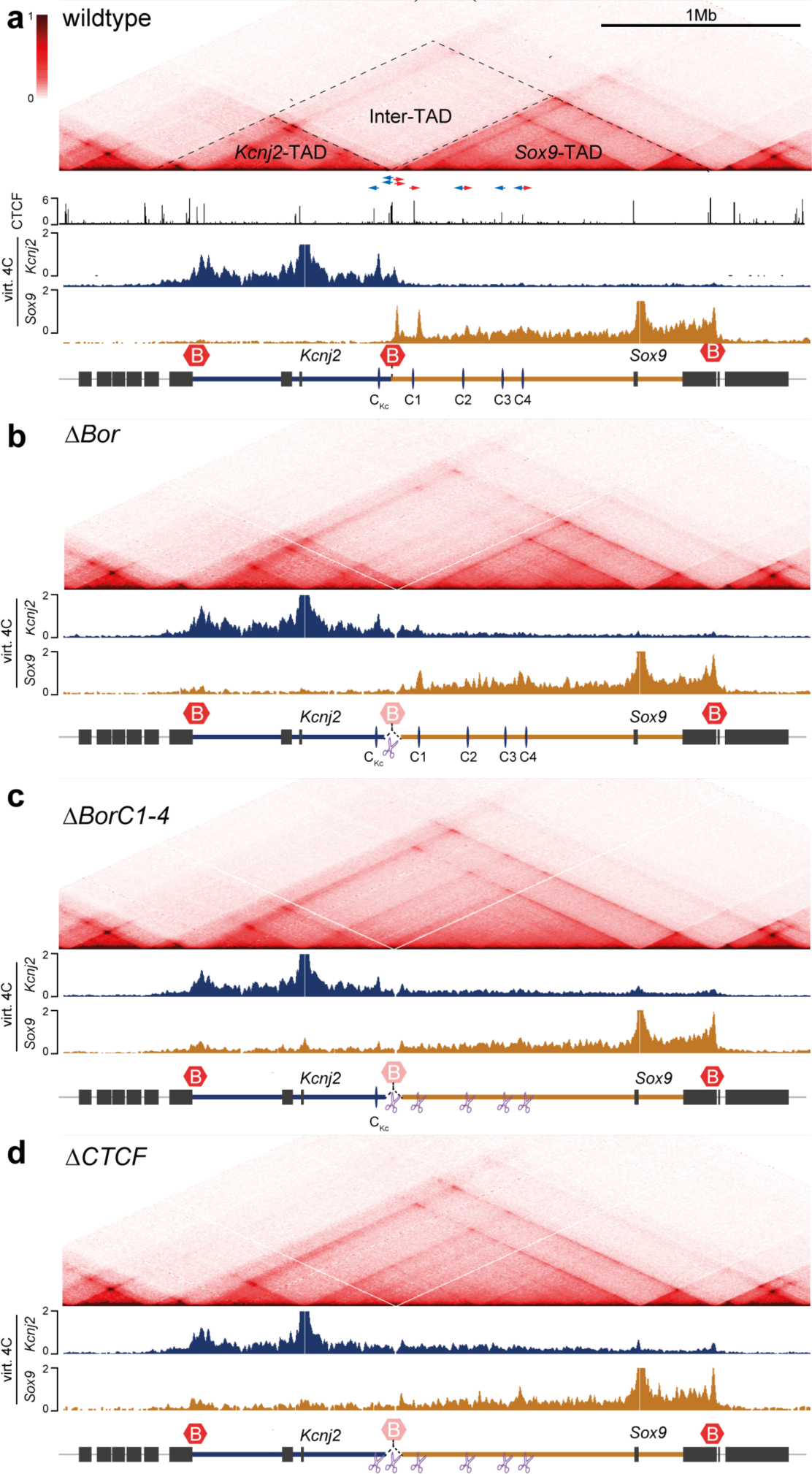
Progressive fusion of the *Kcnj*- and *Sox9*-TADs upon deletion of TAD boundary and intra-TAD CTCF sites. cHiC from E12.5 mouse limb buds. Virtual 4C with viewpoints at the *Kcnj2* (blue) or the *Sox9* promoter (orange) below. CTCF ChIP-seq with binding site orientation (red/blue) at the boundary and intra-TAD are highlighted. Two-headed arrow indicates two oppositely oriented sites (FIMO p<10^-4^) underlying the ChIP-seq peak. (A) Wildtype cHiC. Dashed lines indicate *Kcnj* and *Sox9*-TADs and area of inter-TAD contacts. (B) 18kb-deletion (*ΔBor*) of the TAD boundary leaves TAD configuration largely unchanged. (C) Deletion of the TAD boundary and targeted deletion of all four major intra-TAD CTCF-binding sites (*ΔBorC1-C4*) cause TAD fusion (D) *ΔCTCF* shows further TAD fusion upon deletion of the TAD boundary and all CTCF-binding-sites between the *Kcnj2*- and *Sox9*-promoter (C1-C4 and C_Kc_).

**Figure 3:**
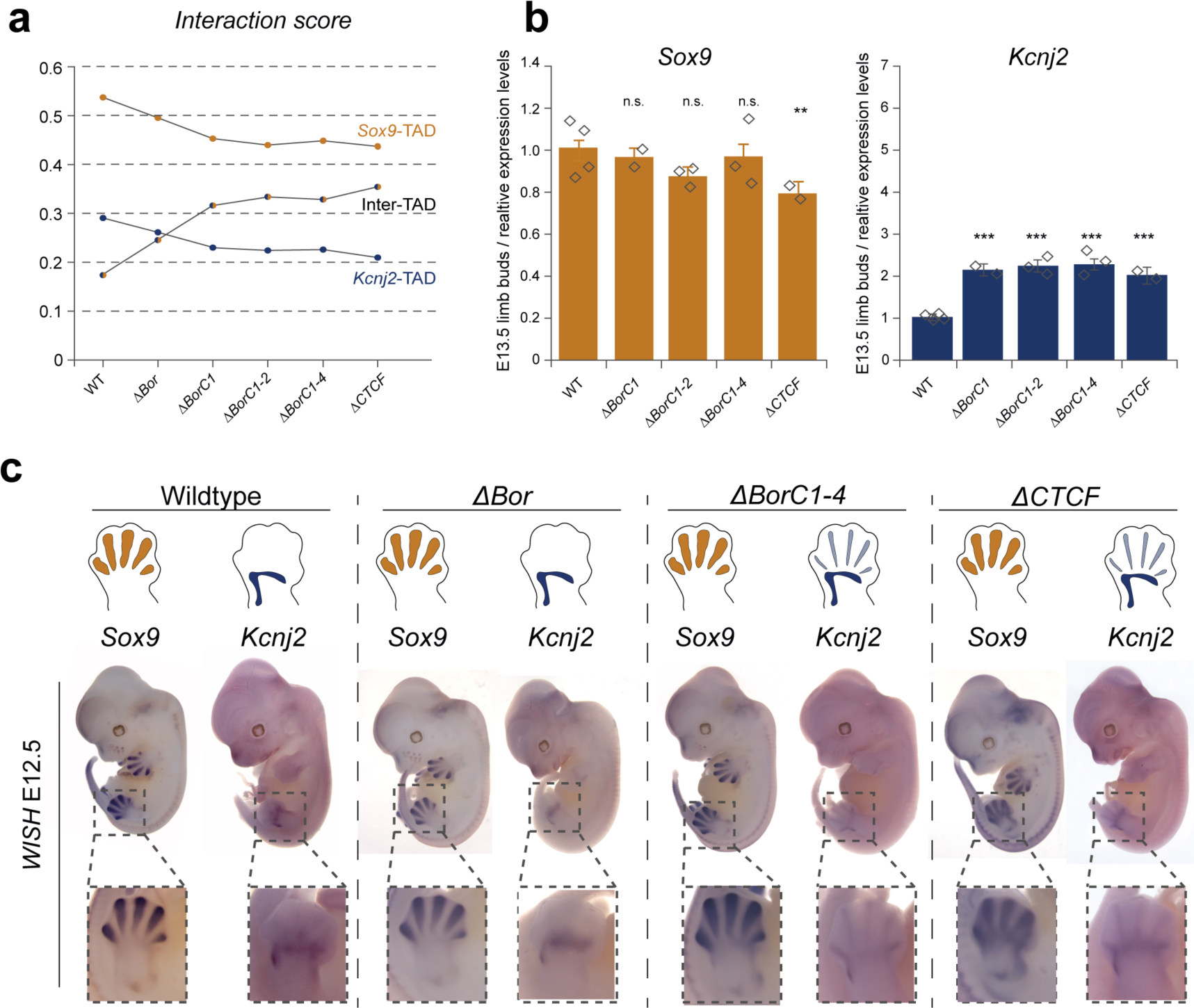
Effect of TAD fusion and deletion of CTCF binding sites on gene expression and phenotype. (A) Change of interaction score induced by consecutive removal of CTCF sites. Percentage of contacts in the *Kcnj2*, and *Sox9*-TADs and between TADs (inter-TAD). The more CTCF-binding sites are deleted, the higher the contact frequency between the *Kcnj2*- and *Sox9*-TAD. (B) Relative gene expression of *Kcnj2* and *Sox9* in E13.5 limb buds measured by real-time qPCR. Values are normalized to *Gapdh*-expression (Wildtype = 1). Bars represent the mean, error bars the standard deviation, diamonds indicate individual replicates. Significance in comparison to wildtype levels tested with unpaired t-test (p<0.01:**, p<0.001:***) (C) Expression pattern of *Sox9* and *Kcnj2* in E12.5 limbs (*WISH*). Schematic on top, whole-mount in situ hybridization below, detailed view of hindlimbs at the bottom. *Sox9* is strongly expressed in the digit anlagen, whereas *Kcnj2* is expressed weakly in the distal stylopod. Note no change in *Sox9* expression and low degree of digit expression of *Kcnj2* only in *ΔBorC1-4* and *ΔCTCF*.

### Loss of TAD insulation has minor effects on developmental gene regulation

We next investigated how the gradual fusion of TADs might affect the expression of *Kcnj2* and *Sox9* and the phenotype. We previously demonstrated that misexpression of *Kcnj2* in a *Sox9*-like pattern results in a malformation of the terminal phalanges (in humans called Cooks syndrome, OMIM 106995) ^15^. Heterozygous loss of *Sox9*, on the other hand, leads to a lethal skeletal phenotype characterized by bowing of long bones, cleft palate, and rib abnormalities in heterozygous animals (in humans called campomelic dysplasia, OMIM 114290) and homozygous mutants do not form cartilage at all ^18^. In addition, *Sox9* is an essential factor for testis development downstream of SRY and its inactivation results in male-to-female sex reversal ^19^. To monitor gene expression changes in embryos we used qPCR and whole-mount in situ hybridization (WISH). Phenotypes associated with loss of *Sox9* and/or gain of *Kcnj2* were assessed in mice by visual inspection (palate, claws), skeletal preparations, µCT (digits) and by testing fertility through breeding.

Despite the observed fusion of TADs, we did not observe any dramatic changes in gene expression. While the changes in *Sox9* gene expression were not significant in the boundary deletion or the *ΔBorC1-4* mutants, we detected a ~10-15% reduction of *Sox9* in the *ΔCTCF* animals (Fig. 3b, Fig. S2a). *Kcnj2*, which is only marginally expressed at this timepoint, increased slightly to approx. 2-fold in all alleles. To detect if there were changes in the patterns of expression in developing limbs, we performed WISH in E12.5 embryos. In all lines, *Sox9* expression was indistinguishable from wildtype embryos. Also, *Kcnj2* stayed unchanged and no *Sox9*-like misexpression was detected in *ΔBor* or *ΔBorC1-2* animals (Fig. 3a, Fig. S2b). However, upon deletion of four or more CTCF sites in addition to the TAD boundary faint *Kcnj2* expression in the digit anlagen was detected (*ΔBorC1-4* and *ΔCTCF*, Fig. 3c). Importantly, despite the fusion of TAD structures, all mutant animals were viable, bred to Mendelian ratios, and had no detectable phenotype. In particular, no abnormality of the digits/claws was observed (Fig. S2c).

Thus, in spite of the observed TAD fusion, *Sox9* and *Kcnj2* expression remained largely unchanged with no phenotypic effect. These results indicated that enhancers were able to efficiently contact their target gene even without CTCF loops. Removal of all CTCF sites, however, led to a spill over of activity from the *Sox9* TAD to the *Kcnj* TAD indicating that TAD genome organization provides a certain degree of robustness and precision to gene expression at this locus.

### Re-organization of 3D chromatin contacts by TAD boundaries and TAD substructure orientation

TAD fusions induced by large structural variations have been reported to cause gene misexpression and disease ^16^, yet the fusion of TADs induced by CTCF site deletion occurred without major gene misexpression. To investigate this discrepancy further, we produced four different types of inversions/insertions: 1) an inversion of the *Sox9* regulatory domain including the TAD boundary (*InvC*), 2) an inversion of the *Sox9* regulatory domain without the boundary (*Inv-Intra*), 3) an insertion of the boundary alone without inverting the regulatory domain (*Bor-KnockIn*), and, finally, 4) an inversion of the *Sox9* regulatroy domain with the boundary removed (*InvCΔBor*) (summarized in Fig. S3). This combination of alleles allowed us to dissect the role of TAD boundaries and substructure orientation in TAD formation and for structural variations.

cHiC from the inversion of the centromeric 1.1 Mb of the *Sox9-*TAD including the TAD boundary (*InvC*) (E12.5 limb buds) showed a fusion of the inverted part of the *Sox9-*TAD with the *Kcnj*-TAD and a separation of the *Sox9* gene and its remaining TAD from the inverted part (Fig. 4a, b). cHiC and virtual 4C showed that *Kcnj2* was now able to contact the C1-C4 CTCF sites in a similar fashion as *Sox9* in the wildtype situation (Fig. 4a, b).

**Figure 4:**
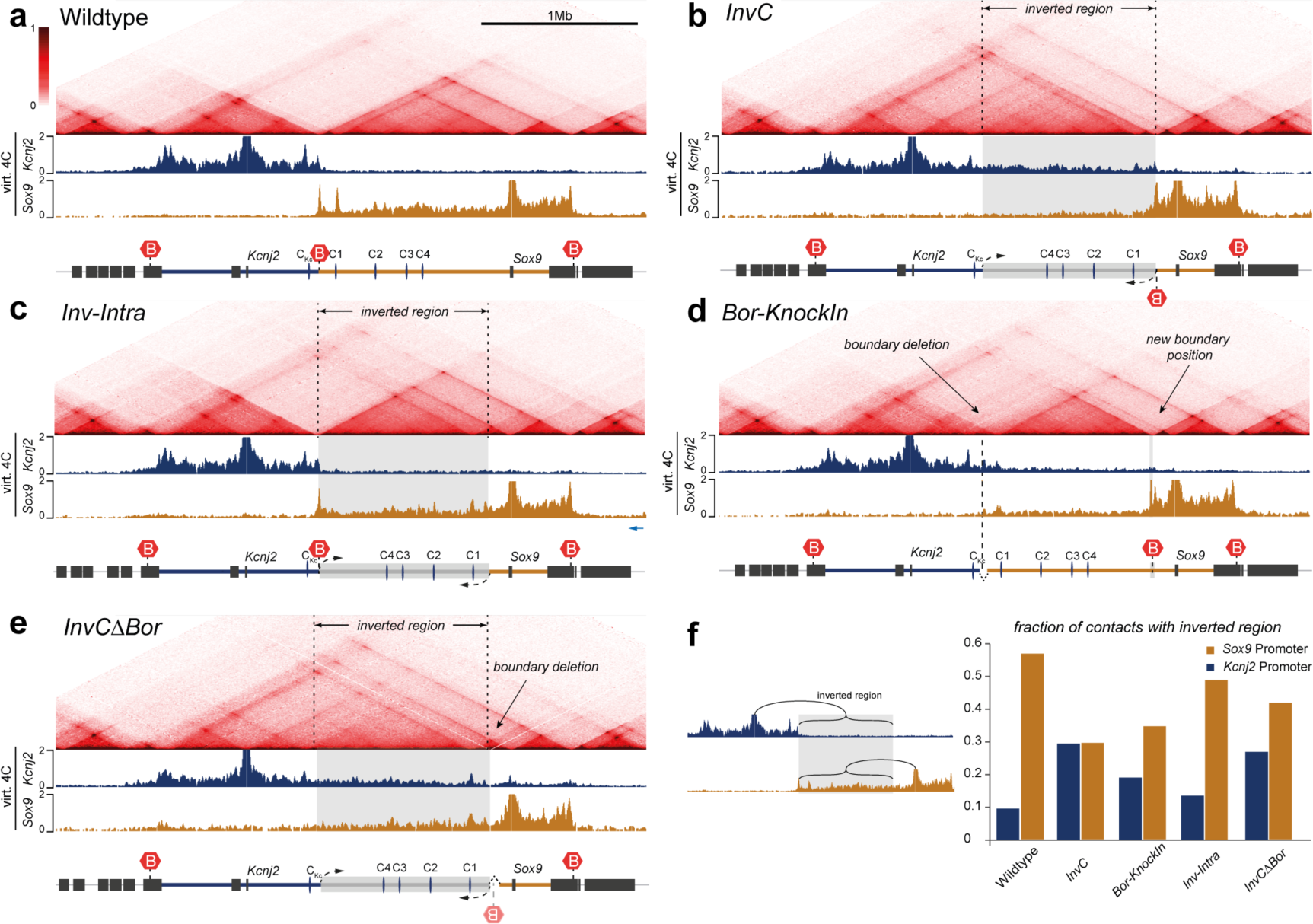
Boundaries and orientation of regulatory landscapes define TAD organization. cHiC from E12.5 mouse limb buds, each mapped to a custom genome. 4C with viewpoint at the *Kcnj2* (blue) or *Sox9*-promoter (orange) below. Grey box indicates the extent of inverted region. (A) Wildtype. (B) Inversion including boundary (*InvC*) leads to fusion of the inverted region with the *Kcnj*-TAD and separation of *Sox9* from its regulatory domain. (C) Inversion excluding the boundary (*Inv-Intra*) has no major changes in TAD configuration. (D) Repositioning of boundary to a new position near *Sox9* (*Bor-KnockIn*) isolates of *Sox9* from its TAD but does not cause the fusion with the *Kcnj*-TAD. (E) InvC inversion with deleted boundary (*InvCΔBor*) causes fusion of both TADs and re-establishes *Sox9* contacts with its TAD. (F) Fraction of contacts with the inverted region for each allele. Bar diagram shows contacts of *Kcnj2* (blue) or the *Sox9* (orange) with the inverted region as a fraction of all contacts in the *Kcnj* and *Sox9* TADs measured by 4C.

To investigate the effect of the inverted TAD substructure vs. the repositioned boundary we produced a slightly smaller inversion not including the boundary (*Inv-Intra*). cHiC of this intra-TAD inversion showed that the overall extent of interactions did not change. However, the re-direction of CGTCF sites resulted in an altered pattern of loop formation (Fig. 4c). The contacts between intra-TAD CTCF sites and the TAD boundary became stronger, while those with the *Sox9* promoter became weaker, as if the entire region had shifted its interaction towards the TAD boundary (Fig. S4).

To test the effect of the TAD boundary alone we used the boundary deletion background to insert a 6.3 kb construct carrying the four boundary CTCF sites (schematic shown in Fig. S5d) 125 kb upstream of *Sox9* (*Bor-KnockIn*). cHiC of this allele showed that the repositioned TAD boundary split the *Sox9*-TAD into two domains. Similar to the TAD-spanning *InvC* inversion, the telomeric region containing the *Sox9* gene was now separated from the centromeric region (Fig. 4d). However, in contrast to the *InvC* inversion, the centromeric *Sox9-*TAD did not fuse with the *Kcnj2*-TAD but remained an isolated domain extending from the C1-CTCF site to the inserted boundary. Thus, the boundary was fully functional even at a different position separating the Sox9 TAD into two domains.

Finally, we wanted to test the effect of TAD substructure orientation without the influence of a nearby boundary. Deletion of the TAD boundary in the *InvC* inversion (*InvCΔBor*) resulted in a loss of insulation and again of *Sox9* contacts with its centromeric TAD. At the same time the contacts between the inverted part of the *Sox9-*TAD and *Kcnj2* were still present (Fig. 4e). Thus, this mutant resulted in a fusion of the entire *Sox9* and *Kcnj*-TADs and both, *Sox9* and *Kcnj2,* were contacting the inverted *Sox9* regulatory region.

To compare how TAD orientation and boundary position affect the contact frequency we quantified the contacts of the *Sox9* and *Kcnj2* promoters with the inverted region (centromeric part of the *Sox9-*TAD) using virtual 4C (Fig. 4f). Analysis of the TAD-spanning inversion (*InvC*) and the boundary knock-in (*Bor-KnockIn*) showed that the repositioned boundary caused a strong reduction of contacts of *Sox9* with its domain, a phenomenon that was not observed when only the TAD substructure was inverted (*Inv-Intra* and *InvCΔBor*). In contrast, inversion of the TAD substructure (*InvC* and *InvCΔBor*) resulted in a gain of contacts of *Kcnj2* with the centromeric *Sox9-*TAD provided that the two regions were not separated by a bondary.

### Re-direction of TAD structure results in distinct regulatory effects and phenotypes

In contrast to the CTCF site deletions, the rearrangements produced pronounced regulatory and phenotypic effects. Animals carrying the *InvC* allele, in which the inverted centromeric *Sox9*-TAD fused with the *Kcnj*-TAD, showed a clear loss-of-function of *Sox9* and a gain-of-function of *Kcnj2*. *Sox9* expression decreased by 20% in heterozygous and 50% in homozygous animals (Fig. 5b, Fig. S5a). As a consequence, heterozygous animals displayed delayed ossification of the skeleton and homozygous animals showed perinatal lethality with all hallmarks of a *Sox9* loss-of-function phenotype (i.e. bowing of long bones, delayed ossification, cleft palate)(Fig. S5c). *Kcnj2* expression increased up to 5-fold in homozygous embryos (Fig. 5b, Fig. S5a). WISH in E12.5 embryos revealed a strong misexpression of *Kcnj2* in a *Sox9*-like pattern in the digit anlagen (Fig. 5c). Heterozygous *InvC* animals showed malformed terminal phalanges with high penetrance, the phenotype associated with *Kcnj2*-misexpression in a *Sox9* like pattern. Homozygous animals had severely dysplastic digits preventing the development of this phenotype. Thus, the TAD-spanning inversion *InvC* reorganized the TADs at the locus and re-directed regulatory activity from *Sox9* to *Kcnj2*.

**Figure 5:**
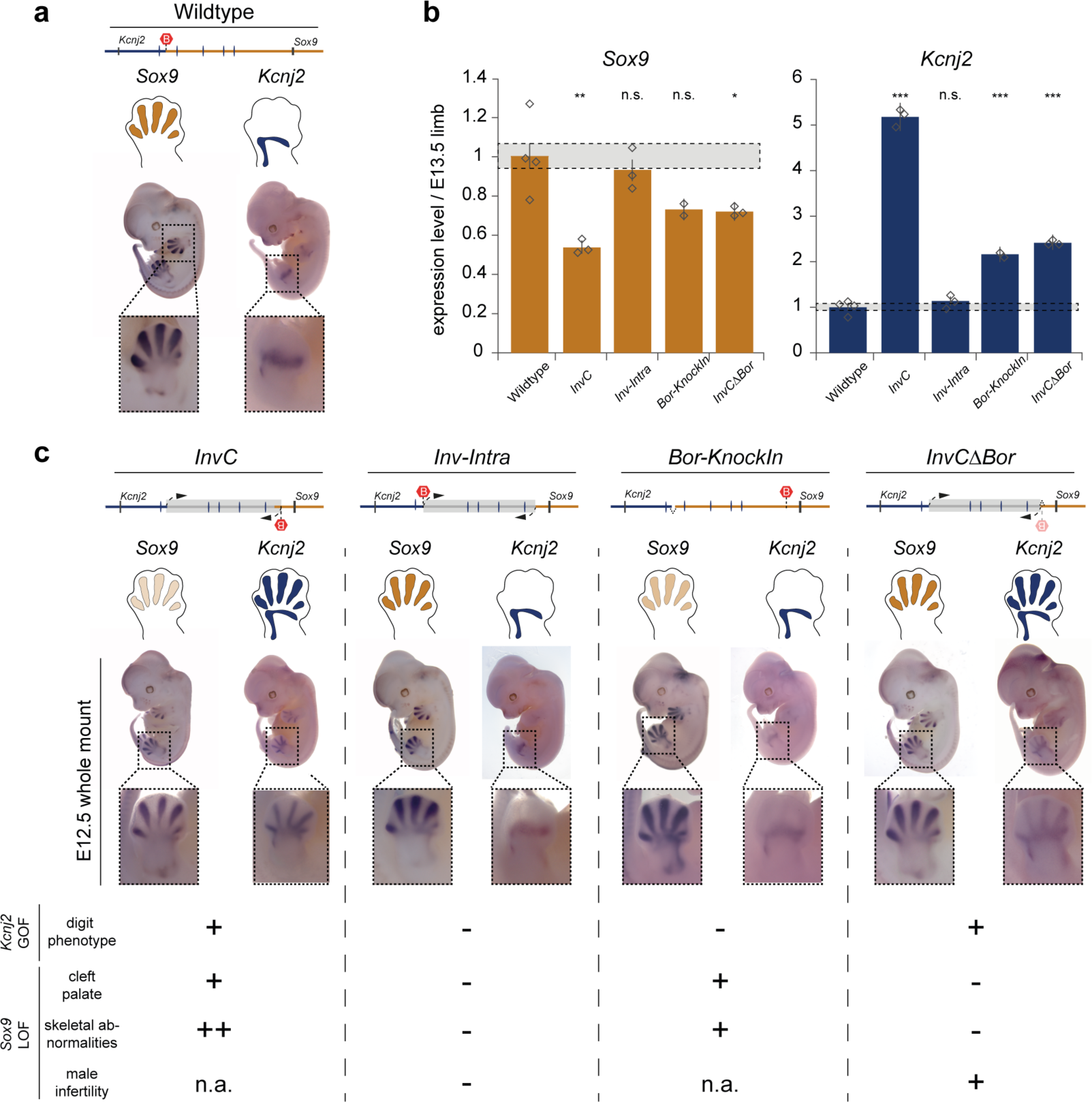
Re-direction of regulatory activity results in *Kcnj2*-misexpression and loss of *Sox9* expression. (A) Wildtype gene expression pattern of *Sox9* and *Kcnj2* (B) Relative gene expression of *Kcnj2* and *Sox9* in E13.5 limb buds measured via real-time qPCR. Values are normalized to *Gapdh*-expression (Wildtype = 1). Bars represent the mean, error bars the standard deviation, diamonds indicate individual replicates. Significance in comparison to wildtype levels tested with unpaired t-test (n.s. not significant, p<0.05:*, p<0.01:**, p<0.001:***) (C) Schematic of Structural Variant. Below, *WISH* of *Sox9* and *Kcnj2* in E12.5 limbs. Schematic on top, whole-mount in situ hybridization below, detailed view of hindlimbs at the bottom. *Kcnj2* gains *Sox9*-like expression pattern in digits in *InvC* and *InvCΔBor*. *Sox9* expression is visibly reduced in *InvC* embryos. Summary of phenotypes in animals induced by gain of *Kcnj2* expression and/or loss of *Sox9* expression.

In contrast, animals with the slightly smaller *Inv-Intra* inversion had no abnormal phenotype. Mice were viable and fertile suggesting no major misregulation of *Sox9* or *Kcnj2*, which was confirmed by qPCR and WISH (Fig. 5b,c). These results indicate that the orientation of the internal TAD structure has no major effect as long as the boundary is intact.

Consistent with the cHiC pattern, the knock-in of the border (*Bor-KnockIn*) showed a *Sox9* loss-of-function, but no *Kcnj2* gain-of-function. *Sox9* levels were reduced by ~40% in homozygous E13.5 limb buds (Fig. 5b) and *Kcnj2* was upregulated ~2-fold, similar to the CTCF-deletion alleles and WISH showed no *Sox9*-like *Kcnj2* misexpression in E12.5 limb buds (Fig. 5a,c). Phenotypically, homozygous *Bor-KnockIn* animals died perinatally due to *Sox9* related defects including cleft palate, short snout, shortened long bones, and delayed ossification (Fig. S5d). However, the skeletal phenotypes were less severe than those seen in the *InvC* animals. Importantly, in accordance with the regulatory effects of the boundary knock-in, the animals had normal phalanges (Fig. S5d).

Finally, we tested gene expression and phenotypes in the *InvCΔBor* allele, in which the entire *Kcnj-* and *Sox9-*TADs are fused and the centromeric *Sox9-TAD* contacted both *Sox9* and *Kcnj2*. Consistent with the re-established cHiC interactions, *Sox9* expression was less severely reduced than in the *InvC* inversion (Fig. 5b, Fig. S5a) and the *Sox9* expression pattern in E12.5 WISH was indistinguishable from wildtype *Sox9* expression (Fig. 5c). On a phenotypic level, the *InvCΔBor* allele rescued the *Sox9* loss-of-function effects of the *InvC* inversion. Homozygous *InvCΔBor* animals had no cleft palate and were viable. The only obvious *Sox9* related phenotype of this allele was that homozygous males were infertile (Fig. 5c). Nonetheless, the *Kcnj2* gain-of-function effects were still present. E13.5 *Kcnj2* expression levels were 2.5-fold higher than wildtype, but lower than in the *InvC* allele (Fig. 5b, Fig. S5a). WISH, however, showed a clear *Sox9*-like expression pattern in the limb buds.

Consistent with this *Kcnj2* misexpression, *InvCΔBor* animals showed the abnormal terminal phalanx phenotype (Fig. S5b).

Thus, phenotype inducing alterations in gene expression can be caused by a change in directionality of the TAD substructure (including their CTCF sites) and/or the repositioning of boundary elements. Such re-directing of regulatory activity can result in either a gain of expression if enhancers are forced to act on a neighboring gene, or in a loss if a gene is disconnected its regulatory domain.

## Discussion

Here we functionally dissect the role of TAD boundaries, intra-TAD CTCF sites and directionality of TAD substructures for TAD formation and gene regulation in a developmental *in vivo* model at the *Kcnj-Sox9* locus. With its two large TADs and the distinct expression patterns of its corresponding genes as well as associated phenotypes, this region is ideally suited for this approach.

### TADs are formed by a redundant system of CTCF sites

The deletion of the *Kcnj-Sox9-*TAD boundary did not have a major effect on the overall TAD configuration. To achieve effective TAD fusion, intra-TAD CTCF sites needed to be deleted in addition to the TAD boundary. Thus, we demonstrate a redundancy in spatial separation of TADs that originates from the combinatorial action of CTCF sites at the TAD boundary and within the TAD. A similar resilience of TAD structures was recently reported for TADs at the HoxD gene cluster, which itself acts as a strong TAD boundary. Here, only a 400kb deletion encompassing the entire HoxD cluster and two flanking genes leads to fusion of the centromeric and telomeric HoxD-TADs ^20^. Smaller deletions do not affect the spatial separation of the two adjacent TADs, supporting our finding that TADs are formed by redundant and buffered mechanisms. Similarly, at the *EphA4* locus the presence of a TAD boundary in pathogenic deletions determines whether two TADs fuse, demonstrating that it functions as a potent insulator. The resulting TAD fusion at the *EphA4* locus leads to misexpression of *Pax3*. This, however, is the result of a deletion that removes not only the boundary but, in addition, the majority of the *Epha4* TAD and alters the entire 3D structure at the locus. In contrast, the serial deletions of CTCF-sites here leave the overall configuration of the locus intact but modify the barrier function between TADs. We thereby provide direct evidence that TADs are formed by a redundant system of CTCF sites at the TAD boundaries and within the TADs.

### Insulation between TADs is not required for developmental gene regulation

We addressed the role of CTCF at an individual locus in an *in vivo* developmental setting and avoid the genome wide effects associated with a loss of CTCF. Surprisingly, the gradual fusion of the two neighboring TADs was accompanied by only mild effects on gene regulation, indicating that the enhancers were able to find, contact, and regulate their cognate promoters. Surprisingly, this loss of TAD structures at the *Sox9* locus also had no phenotypic consequences, indicating that there were no substantial effects on *Sox9* or *Kcnj2* regulation throughout development. These results are in agreement with the weak gene expression changes observed upon CTCF depletion ^8, 14^, suggesting that enhancer-promoter communication can function independent of TADs and CTCF-mediated genome architecture. Such a mechanism might be mediated by homotypic interaction of TFs that bind at distal enhancers and their cognate promoters and could promote transcriptional condensates as recently proposed ^21^. In such a scenario, TADs would act as three-dimensional scaffolds that optimize such interaction hubs, without being essential to establish them.

Another reason for the mild effects on gene regulation is likely that the deletion of CTCF sites does not affect cohesin recruitment to the chromatin, which is independent of CTCF ^22^. Thus, cohesin complexes at the *Sox9* locus can still facilitate enhancer-promoter interaction. What does change, however, are the limits for the extrusion complexes that are normally set by CTCF. The consecutive removal of the boundary and intra-TAD CTCF sites leads to ever increasing contacts of *Kcnj2* with the active Sox9 regulatory landscape. However, this only results in an incremental increase in *Kcnj2* expression indicating that spurious contacts do not directly result in gene regulation.

However, it has to be pointed out that the loss of CTCF/insulation was accompanied a loss of *Sox9* and gain of *Kcnj2* expression induced by a spread of regulatory activity from the *Sox9* to its neighboring TAD. Thus, TADs and their boundaries are probably not essential for developmental gene expression, but they confer precision and robustness to the system. At the locus investigated here, the relatively mild expression changes had no phenotypic effect. However, in other cases where precision and insulation is essential, such leaky expression might result in disease phenotypes.

Our findings support the idea that the spatial separation into TADs and enhancer-promoter interaction represent two independent layers of long range gene regulation. These layers stabilize each other but are not inherently linked. Furthermore, they contradict the generally accepted idea that enhancers are promiscuous ^6^, i.e. can and will activate any promoter in their vicinity. The readiness to become activated by spurious enhancer contacts is likely to depend on various mechanisms including enhancer-promoter specificity ^23^, histone modification, proximity to the target promoter, and openness of chromatin (Kraft et al. 2019).

### Boundaries and the orientation of substructures organize regulatory domains

With our series of inversions and knock-in alleles, we were able to dissect the relevance of TAD substructure and TAD boundaries for altering gene expression. In the TAD-spanning *InvC* inversion the repositioned TAD boundary serves as a strong insulator separating *Sox9* from its regulators. At the same time, re-direction of the TAD substructure creates new loops with the *Kcnj2* gene and fuses the *Sox9* regulatory domain with the *Kcnj2*-TAD. This combination is apparently sufficient to connect the *Sox9* enhancers to the *Kcnj2* promoter, thereby overcoming their inherent affinity for the *Sox9* promoter. As a consequence, we observe a *Sox9* loss of function phenotype and a *Kcnj2* gain of function. Thus, misexpression and disease can be induced by redirecting TAD substructures and enhancer activity but not by removing them. In this context, boundary and TAD substructure function together, but the substructure with its CTCF sites cannot override a boundary. However, its inversion is needed to achieve pathogenic misexpression.

Our results help to explain the apparent discrepancy between the modest effects of CTCF/cohesin depletion on transcription and the drastic effects of TAD reorganization in pathogenic structural variations. Based on our findings it is to be expected that, due to the high redundancy of CTCF sites in maintaining TAD structure, most SVs are likely to be tolerated. To result in aberrant gene activation, the rearrangement needs to actively re-organize 3D chromatin contacts and thereby connect a regulatory region to the new target gene. Such effects can be achieved by deleting boundaries together with adjacent divergent CTCF sites, or by re-directing regulatory activity through inversions or duplications. The re-positioning of boundaries, on the other hand, can result in loss of expression induced by disconnecting a gene from its regulatory domain. Thus, SV induced misexpression is not caused by the simple removal of barriers or the effect single enhancer-promoter rewiring. Rather, it is the result of connecting larger regulatory structures with novel target genes through CTCF mediated loops.

## Methods

### ES cell targeting and transgenic mouse strains

ES cell culture was performed as described previously ^15^. A list of sgRNAs used to generate the various deletions and inversions is given in Sup. Tab. XYZ. For targeting and re-targeting of CTCF-sites, sequence-verified ESC-lines were re-targeted with either one or two pX459-sgRNAs. For each CTCF-site deletion, structural variants were excluded and modified CTCF-sites for both alleles were verified through PCR-amplification of the cut-site, followed by sub-cloning and Sanger-sequencing of several PCR-products. Only if the successful modification of both alleles could be verified, the ESCs were used for further experiments. The results were later validated using the cHi-C sequencing data.Embryos and live animals from ES cells were generated by di- or tetraploid complementation^24^. Genotyping was performed by PCR analysis.

### CRISPR-guided knock-in in mouse ESCs

For targeting the Kcnj-*Sox9*-TAD boundary, a 6.3 kb construct containing the four CTCF sites (C1 site (mm9 chr11:111384818--111385832 followed by C2-4 chr11:111393908-111399229) was cloned into a targeting vector with assymetric homology arms (HA1: chr11:112511756--112514691, HA2: chr11:112514692-12519932) using standard cloning procedures.

For knock-in of targeting constructs without selection marker, the targeting construct was transfected in combination with a pX459-sgRNA vector. Importantly, the targeting construct did not contain either an intact PAM-site or guide-sequence. Puromycin selection and clonal ES cell line generation was performed as described previously. Successfully targeted ESC-lines were screened using PCR and validated for locus-specific integration after successful establishment of the ESC line. Validation of the homozygous TAD-boundary knock-in was performed bioinformatically using the cHi-C data.

### LacZ-Sensor mouse lines

The SB-Kcnj and SB-*Sox9* alleles described in Franke et al. were used for remobilization of the SB transgene, following the protocol in Ruf, et al. ^25^, to generate new SB insertion sites (LacZ-Sensors) at the locus (Kcnj-TAD: SB20, SB16, SB24; *Sox9*-TAD: SB23, SB18). An additional LacZ-Sensors in the *Sox9*-TAD (mid-*Sox9*) was targeted directly using CRISPR-guided knock-in as described above. Asymmetric homology arms (0.8 and 1.5kb) with mutated PAM sites and restriction sites for cloning of the LacZ-transgene were obtained from IDT, cloned into a plasmid vector. The beta-Globin-LacZ-transgene was then inserted into the Acc65I linearized targeting vector using Gibson assembly A list of primers used for cloning and genotyping targeting construct, and pX459-sgRNAs are provided in Supplementary Table 1.

### Generation of mice

Mice from transgenic and genome edited ESCs were generated by di- or tetraploid aggregation (REF), maintained by crossing them with C57Bl.6/J mice, and genotyped by PCR. Primers for genotyping can be provided upon request. All animal procedures were conducted as approved by the local authorities (LAGeSo Berlin) under the license number #G0368/08 and #G0247/13.

### Expression analysis

RNA for qPCR was extracted from E13.5 mouse zeugopods. After dissection, samples were immediately frozen in liquid nitrogen, and individual embryos were genotyped. Tissue was lysed in RLT buffer and a .20 gauge syringe and RNA extraction was performed using the RNeasy Mini Kit (Qiagen) according to manufacturer’s instruction. cDNA-synthesis was performed with SuperScriptIII RT (invitrogen) and polyT-Primers and qPCR was performed on an ABI9700.

### Whole-mount *In Situ* Hybridization

E12.5 embryos were subjected to whole mount *in situ* hybridization using standard procedures. *Sox9* and *Kcnj2* probes were generated by PCR amplification using mouse limb bud cDNA (Supplementary Table 1).

### Micro-Computer Tomography

Autopods of seven week old control and mutant mice were scanned using a Skyscan 1172 X-ray microtomography system (Brucker microCT, Belgium) at 5µm resolution. 3D model reconstruction was done with the Skyscan image analysis software CT-Analyser and CT-volume (Brucker microCT, Belgium).

### cHiC data processing

Raw reads were preprocessed with cutadapt v1.15 ^26^ to trim potential low quality bases (-q 20 -m 25) and potentially remaining sequencing adapters (-a and -A option with Illumina TruSeq adapter sequences according to the cutadapt documentation) at the 3’ ends of the reads. Mapping, filtering, and deduplication of the short reads were performed with the HiCUP pipeline v0.5.10 ^27^ (no size selection, Nofill: 1, Format: Sanger). The pipeline was set up with Bowtie2 v2.2.6 ^28^ for mapping short reads to reference genome mm9. For inversions and the Border-KI allele, reads were also mapped to a customized genome, derived from mm9 based on the genotyping of the mutant ESC lines. Juicer tools 0.7.5 ^29^ was used to generate binned contact maps from valid and unique read pairs with MAPQ≥30 and to normalize contact maps by Knight and Ruiz (KR) matrix balancing ^3, 29, 30^For the generation of contact maps, only reads pairs mapping to the enriched genomic region (chr11:109,010,001-114,878,000) were considered and shifted by the offset of the enriched genomic region (109,010,000 bp). For the import with Juicer tools, we used a custom chrom.sizes files containing only the size of the enriched part of the genome. Afterwards, KR normalized maps were exported at 10kb resolution and coordinates were shifted back to their original values.

Subtraction maps were generated from KR normalized maps, which were normalized in a pair-wise manner before subtraction. To account for differences between two maps in their distance-dependent signal decay, maps were scaled jointly across their sub-diagonals. Therefore, the values of each sub-diagonal of one map were divided by the sum of this sub-diagonal and multiplied by the average of these sums from both maps. Afterwards, the maps were scaled by 10^6^ / total sum. cHiC maps as well as subtractions maps were displayed as heatmaps in which (absolute) values above the 98.5^th^ percentile were truncated for visualization purposes.

### Virtual Capture-C

In order to obtain individual interaction profiles for specific viewpoints with more fine-grained binning, we created virtual Capture-C like interactions profiles from the same filtered bam files that were used for the cHiC maps. Paired-end reads with MAPQ>=30 were considered in a profile when one mate mapped to the viewpoint region, while the other one mapped outside of it. Contacts of the viewpoint region with the rest of the genome were counted per restriction fragment. Afterwards, count data was binned to 1 kb bins. In case a restriction fragment overlapped with more than one bin, the counts were split proportionally. Afterwards, each profile was smoothed by averaging within a sliding window of 5 kb and scaled by 10^3^ / sum of its counts within the enriched region. The viewpoint and a window ±5kb around it were excluded from the computation of the scaling factor. The profiles were generated with custom Java code using htsjdk v2.12.0 (https://samtools.github.io/htsjdk/).

### Interaction score between Kcnj2 TAD and Sox9 TAD

The Kcnj2 TAD was manually defined as genomic region chr11:110,340,001-111,400,000 and the Sox9 TAD as chr11:111,400,001-113,030,000. Contact counts were summed within each TAD individually and within the region of the cHiC map containing the contacts between the two TADs. To avoid a strong influence of the main diagonal, only contacts spanning more than 100 kb were considered in this analysis. The three sums of contact counts were normalized to represent fractions adding up to 1. Thus, the change of contact frequency between the two TADs was determined with respect to the intra-TAD contact frequency.

For the calculation of the *Sox9*/*Kcnj2* contacts with the centromeric *Sox9* TAD, the virtual 4C-seq interaction data were further processed. The contacts with the “inverted region” represent the respective proportion of all *Sox9*/*Kcnj2* promoter contacts in the *Sox9* and *Kcnj2* TADs. The contacts <50 kb from the promoter were excluded from the calculation.

## Acknowledgements

This study was supported by grants from the Deutsche Forschungsgemeinschaft to SM. We thank Judith Fiedler and Karol Macura (transgenic unit), Myriam Hochradel (sequencing core facility) of the MPIMG; Asita Stiege and Ute Fischer for help with cloning and cell culture and Norbert Brieske for in situ hybridizations. We also thank all members of the Mundlos lab for input and discussions.

## Author Contributions

D.M.I. and S.M., conceived the project. A.D., D.M.I., and M.F. performed cHi-C, R.S. performed the computational analysis with input from M.V. and D.M.I.. D.M.I., A.D., S.A., and M.F. produced transgenics, carried out transgenic validation, and performed expression/phenotypic analysis. C.P. performed ATAC-seq, A.D., I.J. and D.M.I. ChIP-seq. B.T. oversaw sequencing of the cHi-C, ChIP-, and ATAC-seq. L.W. performed morula aggregation. W-L.C. performed micro-CT analysis. D.M.I., A.D., and S.M. wrote the manuscript with input from the authors.

## Data availability

Datasets are available through the Gene Expression Omnibus (GEO) under accession number GSE78109 and GSE125294. Secure token for reviewers: szudooiinjwfpej

## Author Information

The authors declare no competing financial interests. Correspondence and requests for materials should be addressed to S.M. and D.M.I.

---

**Figure S1:**
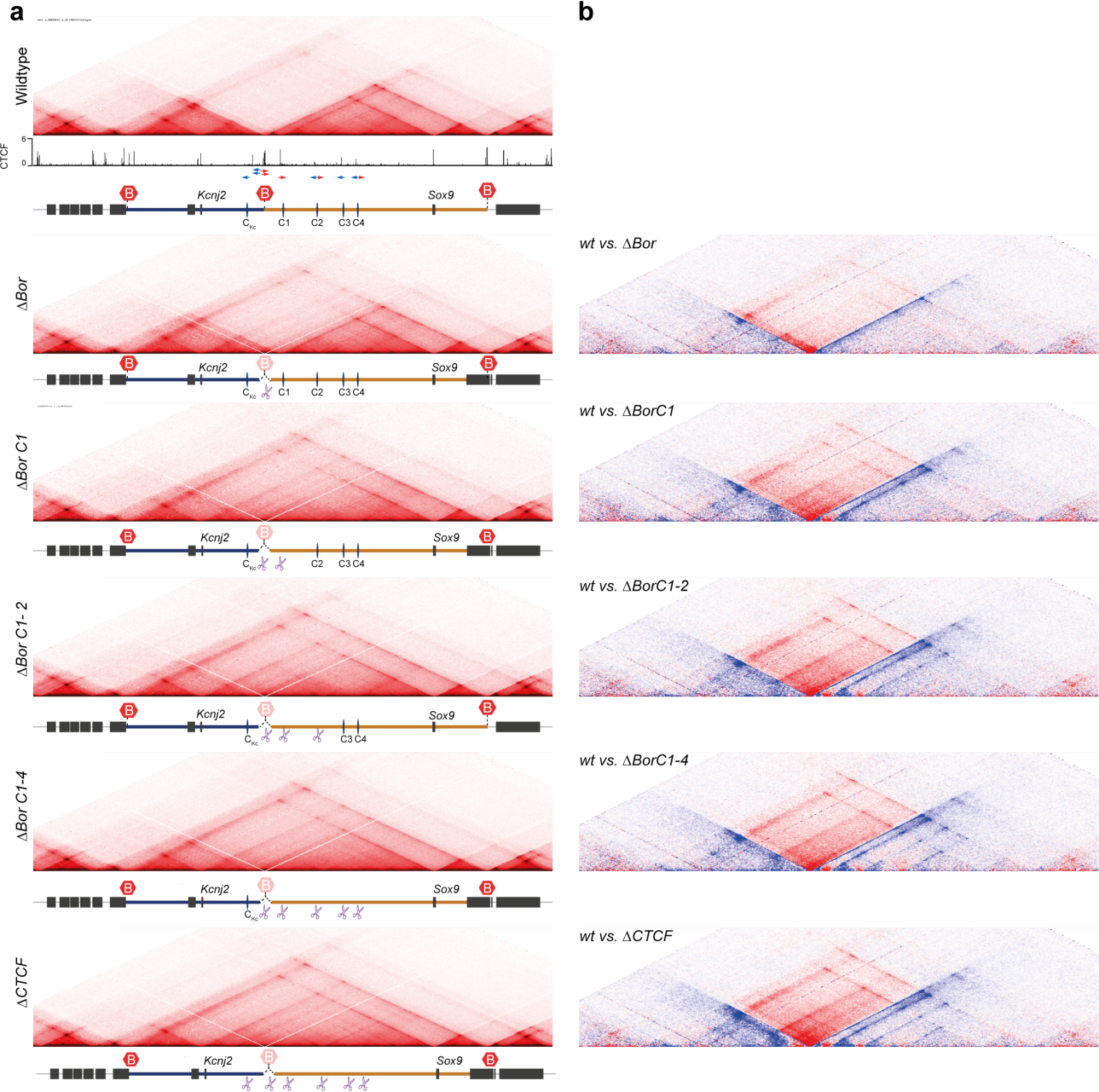
Gradual fusion upon progressive CTCF site deletion. (A) cHiC chromatin interactions from wildtype and mutant E12.5 mouse embryonic limb buds. CTCF ChIP-seq with binding site orientation (red/blue) at the boundary and intra-TAD are highlighted. Two-headed arrow indicates two significant motifs (FIMO, p<10^-4^) underlying the ChIP-seq peak. Virtual 4C from *Kcnj2* (blue) and *Sox9* (orange) below each map. Progressive deletion of CTCF sites causes progressively increasing interaction between *Sox9*- and *Kcnj*-TADs (B) Subtraction maps of wildtype vs. mutant for the same region show increasingly more (red) interactions in the mutants and loss (blue) of the intra-TAD CTCF-mediated contacts.

**Figure S2:**
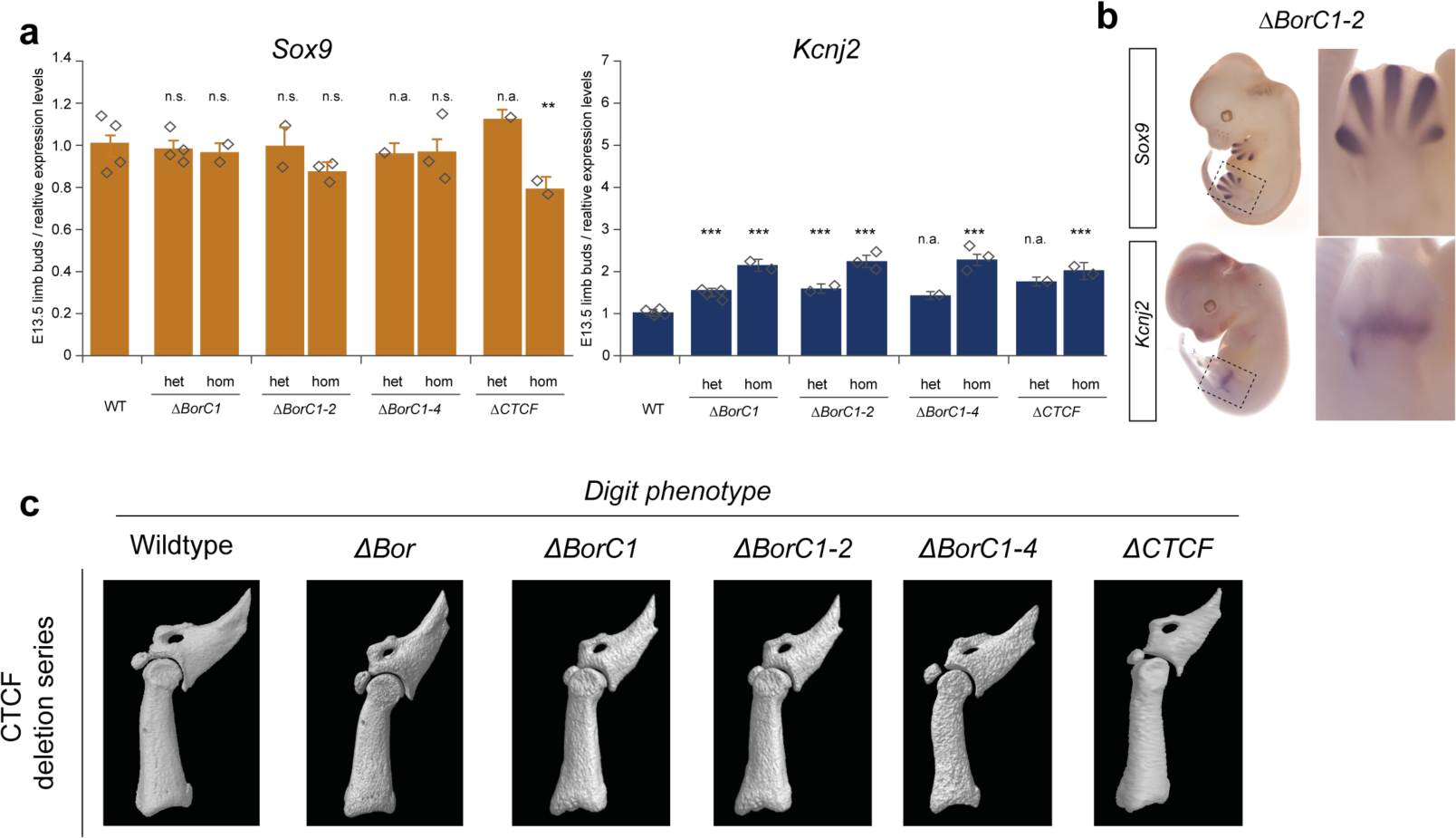
Minor gene expression changes and absence of digit phenotypes upon CTCF-site deletion. (A) *Gapdh* normalized expression of *Kcnj2* and *Sox9* in E13.5 limb buds from heterozygous and homozygous littermates measured via qRT-PCR (wildtype = 1). Bars represent the mean, error bars the standard deviation. Significance relative to wildtype levels tested via unpaired t-test (n.s.: not significant, p<0.05:*, p<0.01:**, p<0.001:***, n.a.: not available) (B) Normal *Kcnj2* and *Sox9* expression pattern in ΔBorC1-2 E12.5 embryos by WISH. Zoom-In shows hindlimb in detail (C) 3D micro-computed tomography scan of terminal phalanges from wildtype and mutant adults (7-12w).

**Figure S3:**
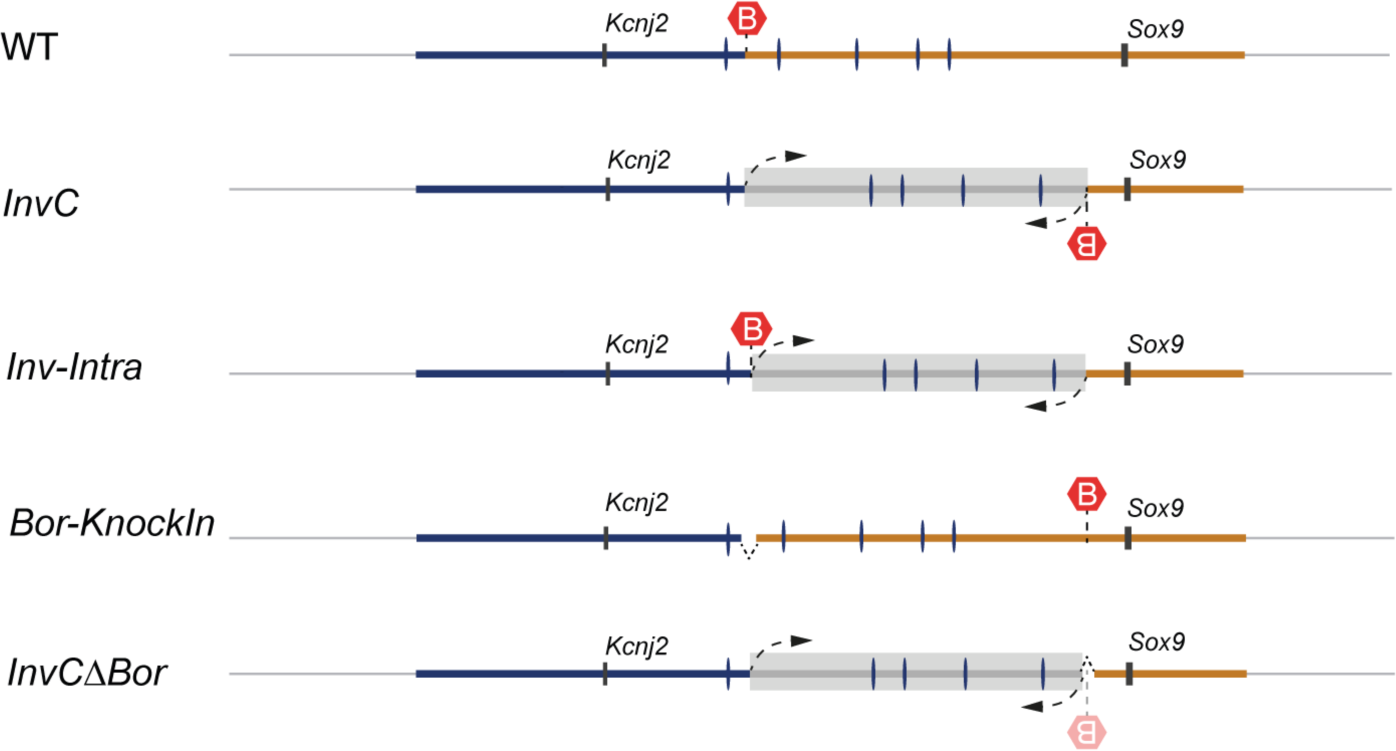
Schematic of the inversion series at the *Kcnj-Sox9* locus. Position of the *Kcnj2* and *Sox9* genes, TAD boundary (red hexagons), and CTCF sites (blue ticks) in wildtype and mutant genomes are indicated. Grey boxes indicate the extent of the inverted region.

**Figure S4:**
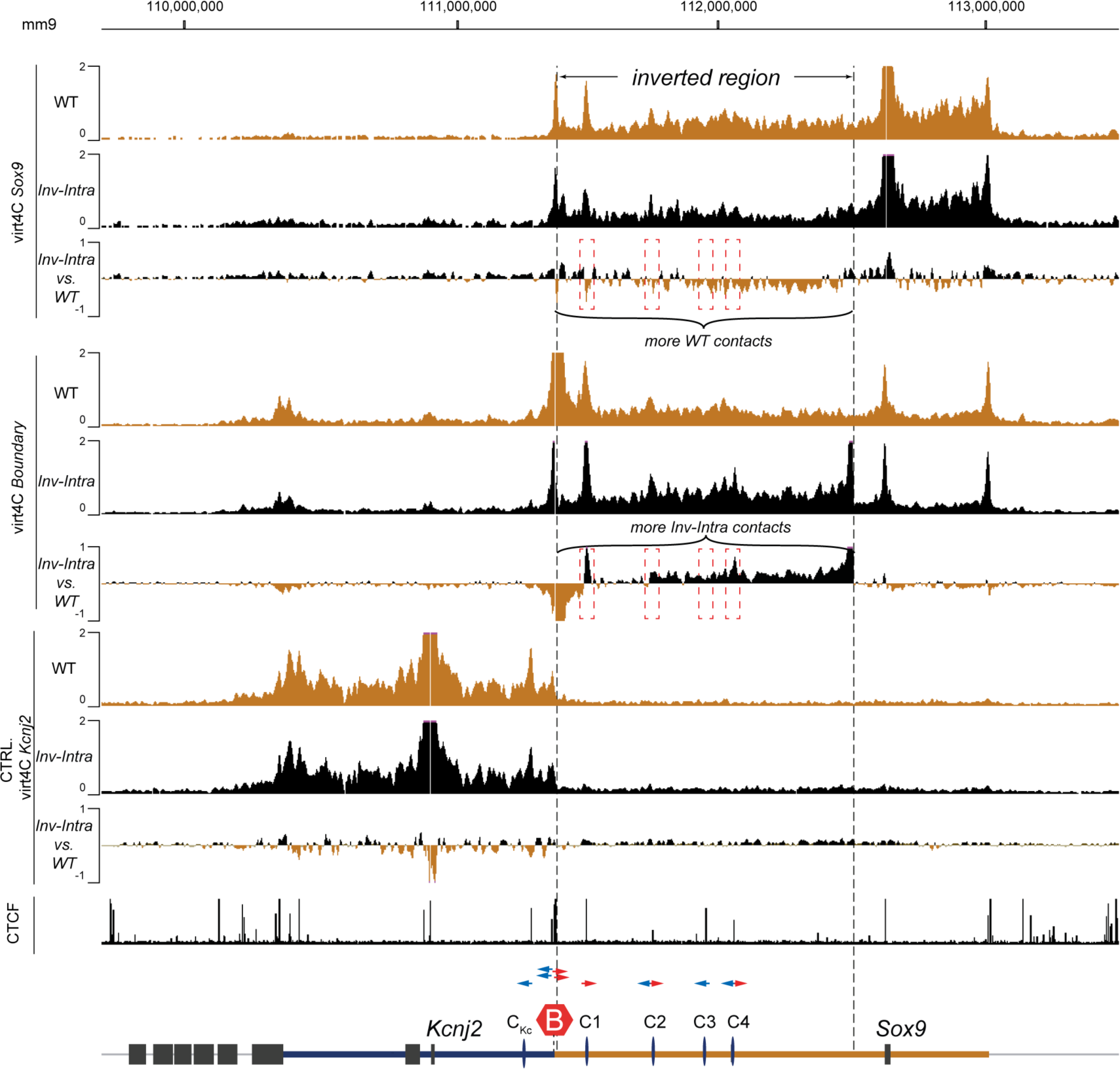
Subtle differences in interactions upon intra-TAD inversion (*Inv-Intra)*. Virtual 4C-profiles derived from cHi-C maps (wildtype and *Inv-Intra*) mapped to a wildtype genome. 4C-profiles from viewpoints at *Sox9*, the TAD boundary, and *Kcnj2* are shown for wildtype (orange) and *Inv-Intra* (orange). Subtraction of the two profiles shows systematically more contacts at the Boundary and *Sox9* viewpoints depending on orientation of TAD substructure for the. The control viewpoint (*Kcnj2*) shows no systematic changes. Note changes in interaction frequency of *Sox9* or the TAD boundary upon inversion of the TAD substructure. Dashed boxes indicate changes in interaction at the C1-C4 CTCF sites.

**Figure S5:**
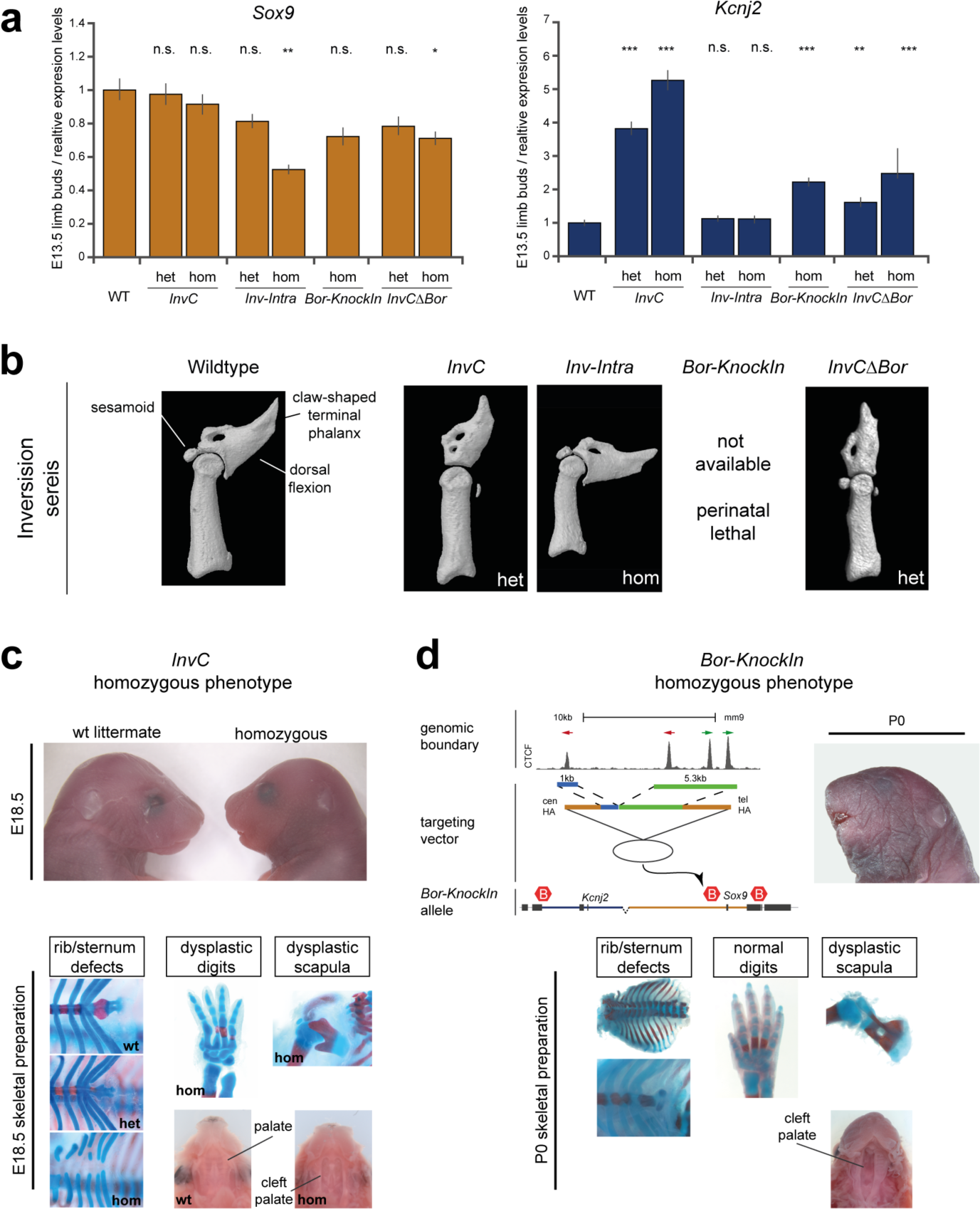
Re-direction of genomic interactions leads to *Sox9* loss and *Kcnj2* gain in expression and developmental phenotypes. (A) *Gapdh* normalized expression of *Kcnj2* and *Sox9* in E13.5 limb buds from heterozygous and homozygous littermates measured via qRT-PCR (wildtype = 1). Error bars represent the std, deviation the mean, diamonds the individual data points. Significance relative to wildtype (unpaired t-test; n.s.: not significant, p<0.05:*, p<0.01:**, p<0.001:***). Note the allele-specific regulatory changes in *InvC* and *InvCΔBor* littermates. Homozygous *Bor-KnockIn* embryos were generated by tetraploid aggregation. (B) Loss of dorsal flexion, sesamoid bones, and claw-shaped form of the terminal phalanx in adult (7-12w) *InvC* and *InvCΔBor* animals shown by 3D micro-computed tomography. (C) Homozygous *InvC* phenotype in E18.5 littermates. Top, lateral view of the head, note short snout and micrognathy in the homozygous embryo. Bottom, Skeletal preparation of littermates show *Sox9*-LOF characteristic skeletal defects (sternum/rib defects, delayed ossification, dysplastic digits and scapula, cleft palate). (D) Homozygous *Bor-KnockIn* phenotype in P0-animals derived from tetraploid aggregation. Top, schematic of the targeting construct containing the four CTCF sites of the TAD boundary (6.3 kb). Homology arms are 2 and 4 kb. Top right: lateral view of P0 newborn, note short snout and micrognathy. Bottom: *Bor-KnockIn* phenotype displays *Sox9*-LOF like defects. Note the overall milder defects compared to *InvC* and absence of a digit phenotype.

